# Auditory Brainstem Responses in Two Closely Related *Peromyscus* Species (*Peromyscus leucopus* and *Peromyscus maniculatus*)

**DOI:** 10.1101/2024.12.09.627419

**Authors:** Luberson Joseph, Emily Margaret New, Desi Marie Joseph, Tamara Chenell Woodley, Vanessa Yamileth Franco, Ben-Zheng Li, Guinevere OU Wogan, Elizabeth A. McCullagh

## Abstract

The genus *Peromyscus* has been extensively used as a model for ecological, behavioral, and evolutionary studies. We used auditory brainstem responses (ABRs), craniofacial morphology, and pinna measurements to compare characteristics that could impact the threshold, amplitude, and latency of ABRs in two wild-caught species, *P. leucopus* and *P. maniculatus*. We observed significant differences in craniofacial and pinna attributes between species, with *P. leucopus* exhibiting larger features and bigger overall size compared to *P. maniculatus*. ABR recordings showed similar hearing thresholds with peak sensitivity between 8 to 46 kHz for both species. We found that amplitude of ABR wave I and IV increased significantly with increasing intensity, while no main effect of species was detected. Latency of wave I and IV significantly decreased with increasing intensity in both species. Finally, we observed longer latencies of the binaural interaction component (BIC) at longer interaural time differences (ITDs) in *P. leucopus*, while no differences were observed across relative BIC amplitude between species. These results provide additional ABR related data that expands the use of both *Peromyscus* species as future models for auditory research.

## INTRODUCTION

Auditory perception and sound localization are essential abilities for survival and fitness across diverse taxa. In small mammals, these abilities facilitate predator avoidance, capturing prey, finding mates, foraging, and conspecific communication (Colburn et al., 1987; Grothe et al., 2010; Kidd et al., 1995). To perceive sound source location, mammals rely on interaural time differences (ITDs) and interaural level differences (ILDs) between the two pinnae. ITD and ILD cues are influenced by the size of the head and the shape of the pinna (Blauert, 1997; Grothe et al., 2010). The auditory brainstem consists of specialized regions that integrate ITD and ILD information from the two ears. Despite decades of research on auditory perception and sound localization in rodent model species (Blauert, 1997; Grothe et al., 2010; Heffner, 2001), our understanding of species-specific biological variation in sound localization and hearing continues to need to be explored (Capshaw et al., 2023).

To understand the mechanisms of hearing, animal models, including laboratory and wild rodents, can serve as valuable tools. Most studies have used the laboratory house mouse (*Mus musculus*) as a model in hearing research due to its sensitive hearing, breeding success and maintanence in laboratory settings, and genetic manipulability (Capshaw et al., 2022; Ehret and Dreyer, 1984). Yet, the limited genomic diversity in inbred laboratory rodents pose challenges to fully recapitulating the broad spectrum of human disorder phenotypes (Voelkl et al., 2020). The house mouse has faced criticism as a model for auditory research, owing to its poor sensitivity to low frequency sounds, increased vulnerability to noise, short lifespan, and audiometric variation within strains (Capshaw et al., 2022, Dammann, 2017). However, the auditory field has leveraged many alternative species including the Mongolian gerbil (*Meriones unguiculatus*), which is valued for its similarity of hearing range to humans (Heeringa et al., 2020; Jüchter et al., 2022; Mills et al., 1990). Although these model taxa have shed insights in understanding the fundamental mechanism of hearing, it is essential to continue to consider comparative approaches that include a wider range of species, particularly those that reflect natural diversity in their vocalizations, have long life spans, and vary in habitat use (Capshaw *et al* 2023).

The white-footed mouse (*Peromyscus leucopus*) and the North American deer mouse (*Peromyscus maniculatus*) are two of the most abundant rodents in North America (Kirkland and Layne, 1989). They both belong to the family Cricetidae and are more closely related to hamsters and voles than to mice of the family Muridae. Both species have been extensively studied as model systems in behavioral, biogeographical, developmental, ecological and evolutionary investigations with regard to their physiological adaptation to varying habitat types (arboreal habitats, grassland, woodlands, brushlands, swamps, and desert), their social system (mainly promiscuous), and behaviors (maternal, winter nesting, climbing, and agonistic behaviors) (Bedford and Hoekstra, 2015; Harney and Dueser, 1987; Lewarch and Hoekstra, 2018). Both species are also of human health concern with regards to their carrying viruses and pathogens including hantavirus, leptospirosis, and plague (Childs et al., 1994; Larson et al., 2018).

The two species differ in their body mass with *P. leucopus* displaying larger body mass and pinna (Light et al., 2021; Kamler et al., 1998). The average body mass of *P. leucopus* was reported to be 26.4 (± 3.9) and that of *P. maniculatus* was 14.5 (± 2.6) (Kamler et al., 1998). *Peromyscus* species hear best between 8 to 16 kHz, as demonstrated by low behavioral and ABR thresholds, with the ability to hear up to 65 kHz (Capshaw et al., 2022; Dice and Barto, 1952; Ralls, 1967). Differences in size may contribute to differences in hearing since pinna size impacts hearing by enhancing sound collection and amplification, improving frequency discrimination, and facilitating more accurate sound localization (Heffner et al., 2020; Heffner and Heffner, 1982).

Recently, members of the genus *Peromyscus* have emerged as valuable model systems in the field of neuroscience for studying age-related hearing loss due to their considerable lifespan compared to mammals of similar size (Capshaw et al 2022). *Peromyscus* species exhibit an average lifespan nearly double that of *M. musculus* when reared in laboratory settings, with the potential to live up to eight years (Burger and Gochfeld, 1992; Guo et al., 1993). In comparison to *M. musculus*, *Peromyscus* rodents display lower production of reactive oxygen species and enhanced resistance to oxidative stress, resulting in delayed accumulation of oxidative damage to deoxyribonucleic acid over the *Peromyscus* lifespan (Csiszar et al., 2007; Labinskyy et al., 2009; Shi et al., 2013), among other preventive effects which may slow cochlear aging (Ohlemiller and Frisina, 2008). Therefore, members of the genus *Peromyscus* show promise as models that can be used to complement auditory research across species and consequently can be reference taxa to explore small mammals’ hearing across longer lifespans.

The purpose of this investigation is to compare hearing-related anatomy, hearing range, monaural and binaural hearing of *P. leucopus* and *P. maniculatus* measured by craniofacial features, pinna size, and auditory brainstem responses (ABRs). We expect that *P. maniculatus* will have shorter binaural latency compared to *P. leucopus*, due to smaller overall body mass, pinna, and craniofacial measurements. We also expect that there will be differences in other measures of hearing including frequency thresholds and monaural ABRs (monaural amplitude and monaural latency) due to differences in body mass, craniofacial features, and other variability between these two closely related species.

## MATERIALS AND METHODS

All procedures used for all experiments complied with the guidelines of the American Society of Mammologists (Sikes and the Animal Care and Use Committee of the American Society of Mammalogists, 2016), were approved by the Oklahoma State University Institutional Animal Care and Use Committee (IACUC), and were conducted with permission from the Oklahoma Department of Wildlife Conservation.

### Animals

Experiments were conducted on 26 individuals, including 15 wild *P. leucopus* (9 males, 6 females) and 11 wild *P. maniculatus* (9 males, 2 females). Animals were live trapped using aluminum Sherman (H.B Sherman Traps, Inc. Tallahassee, FL) non-folding traps (3” x 3” x 10”) between June 2021 and July 2022 at three different locations across Oklahoma, USA (Packsaddle Wildlife Management Area, James Collins Wildlife Management Area, and Payne County) (Figure 1). The traps were baited with old fashioned oats and creamy peanut butter, left overnight, and collected the next morning (∼12 hours). Upon capture, animals were aged based on size, morphologically identified to species in the wild according to (Caire, 1989) and confirmed with Deoxyribonucleic acid (DNA) barcoding polymerase chain reaction (PCR) of collected tail snips. Age was estimated for each animal based on body mass and were divided into three age groups (juvenile, subadult, and adult) (see Table 1). Animals were then transported to the laboratory for ABRs.

**Figure 1:**
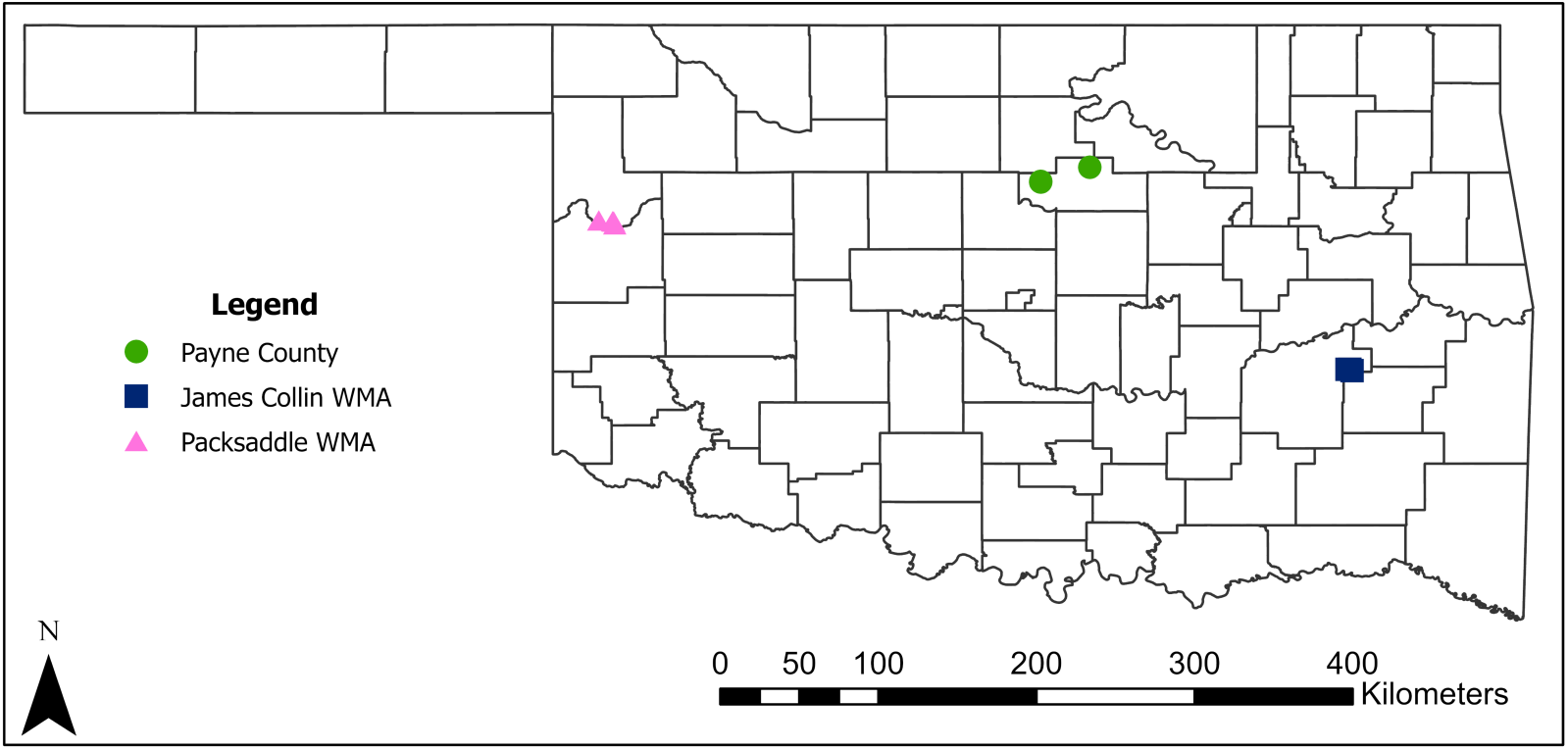
Map showing trapping site locations in Oklahoma. Packsaddle wildlife management area (WMA) sites are represented by pink triangles, James Collin wildlife management area (WMA) sites are represented by blue squares, and Payne County sites are represented by green circles.

**Table 1:**
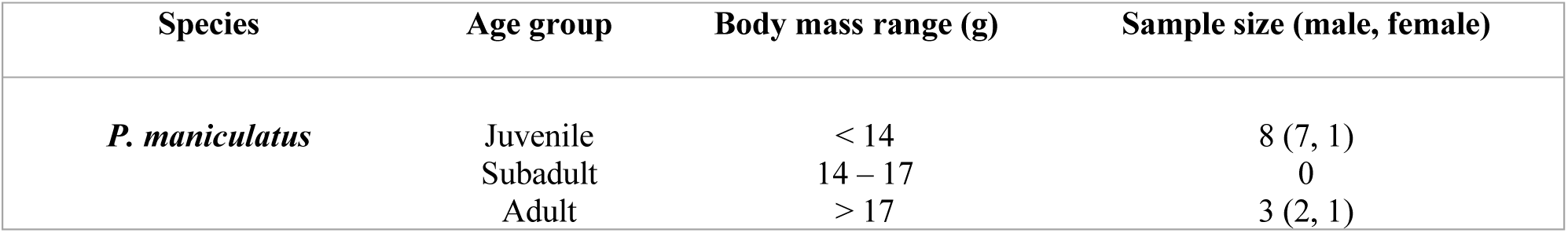
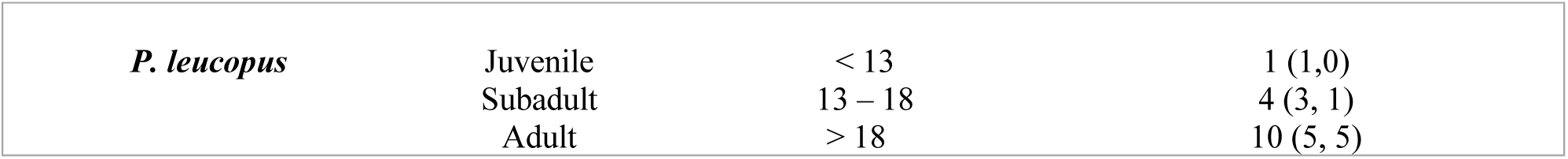
Age was estimated based on body mass for each species based on published literature. Ages for *P. maniculatus* were described as follows: Juveniles < 14 g, subadults, between 14-17 g, and adults, > 17 g (Fairbairn, 1977). We inferred ages for *P. leucopus* as follow: Juveniles < 13 g, subadults, between 13 – 18 g, and adults > 18 g (Cummings and Vessey, 1994). We did not make comparisons by age due to limited sample sizes of each age group.

| Species | Age group | Body mass range (g) | Sample size (male, female) |
| --- | --- | --- | --- |
| *P. maniculatus* | Juvenile | < 14 | 8 (7, 1) |
|  | Subadult | 14 – 17 | 0 |
|  | Adult | > 17 | 3 (2, 1) |
| *P. leucopus* | Juvenile | < 13 | 1 (1,0) |
|  | Subadult | 13 – 18 | 4 (3, 1) |
|  | Adult | > 18 | 10 (5, 5) |

### DNA extraction, amplification, and sequencing

DNA was extracted from tail tissue samples by proteinase K digestion using a Qiagen DNeasy blood and tissue kit (Hilden, Germany) and the protocol outlined by (Nicolas et al., 2012). The DNA concentration and purity were first determined by using a Thermo Scientific Nano-Drop Lite-Spectrophotometer (Fisher Scientific, Spectrophotometer, Nanodrop Lite 6V 18W, Wilmington, DE). The Cytochrome c Oxidase Subunit I (COI) gene was amplified using the primer sequences (CCTACTCRGCCATTTTACCTATG) and (ACTTCTGGGTGCCAAAGAATCA) (Ducroz et al., 2001; Robins et al., 2007). DNA samples were assayed in a 50 μl reaction with 25 μl Phusion Master mix, 2.5 μl forward primer, and 2.5 μl reverse primer and mQ water. The PCR comprised of 35 cycles: 95° C for 300 seconds; 30 seconds at 94° C, 40 seconds at 55° C, 90 seconds at 72° C, and a final 300 second extension at 72° C. The double-stranded PCR products were purified and sequenced at the Center for Genomics and Proteomics of Oklahoma State University (Stillwater, Oklahoma, USA). All sequences were compared with other COI sequences using the NCBI GenBank (Sayers et al., 2022, 2021) databases to confirm species identification (supplemental Figure 1, supplemental Table 1).

### Morphological measures

Craniofacial morphology features including pinna size, tail length, body length, and body mass were recorded for each animal using a six-Inch Stainless Steel Electronic Vernier Caliper (DIGI-Science Accumatic digital Caliper Gyros Precision Tools Monsey, New York, USA) and a digital stainless Steel Electronic scale (Weighmax W-2809 90 LB X 0.1 OZ Durable Stainless Steel Digital Postal scale, Chino, California, USA). Measurements of animal head and pinna including inter-pinna distance (mm) (measurement between the two ear canals), nose to pinna distance (mm) (measurement from the tip of the nose to the middle of the pinna), pinna length (mm) (basal notch to tip, excluding hairs), and pinna width (mm) were measured. Pinna measurements (pinna width and pinna length) were used to calculate the effective pinna diameter which is the square root of the pinna length multiplied by the pinna width (Anbuhl et al., 2017). Tail length (sacrum to caudal tip, excluding hairs), body length (tip of nose to caudal tip), and weight to the nearest gram were taken for each animal (Figure 2A). To assess dependence of morphological traits on body size, log values of traits (pinna width, length etc.) were plotted against the log body length (supplemental Figure 2).

**Figure 2:**
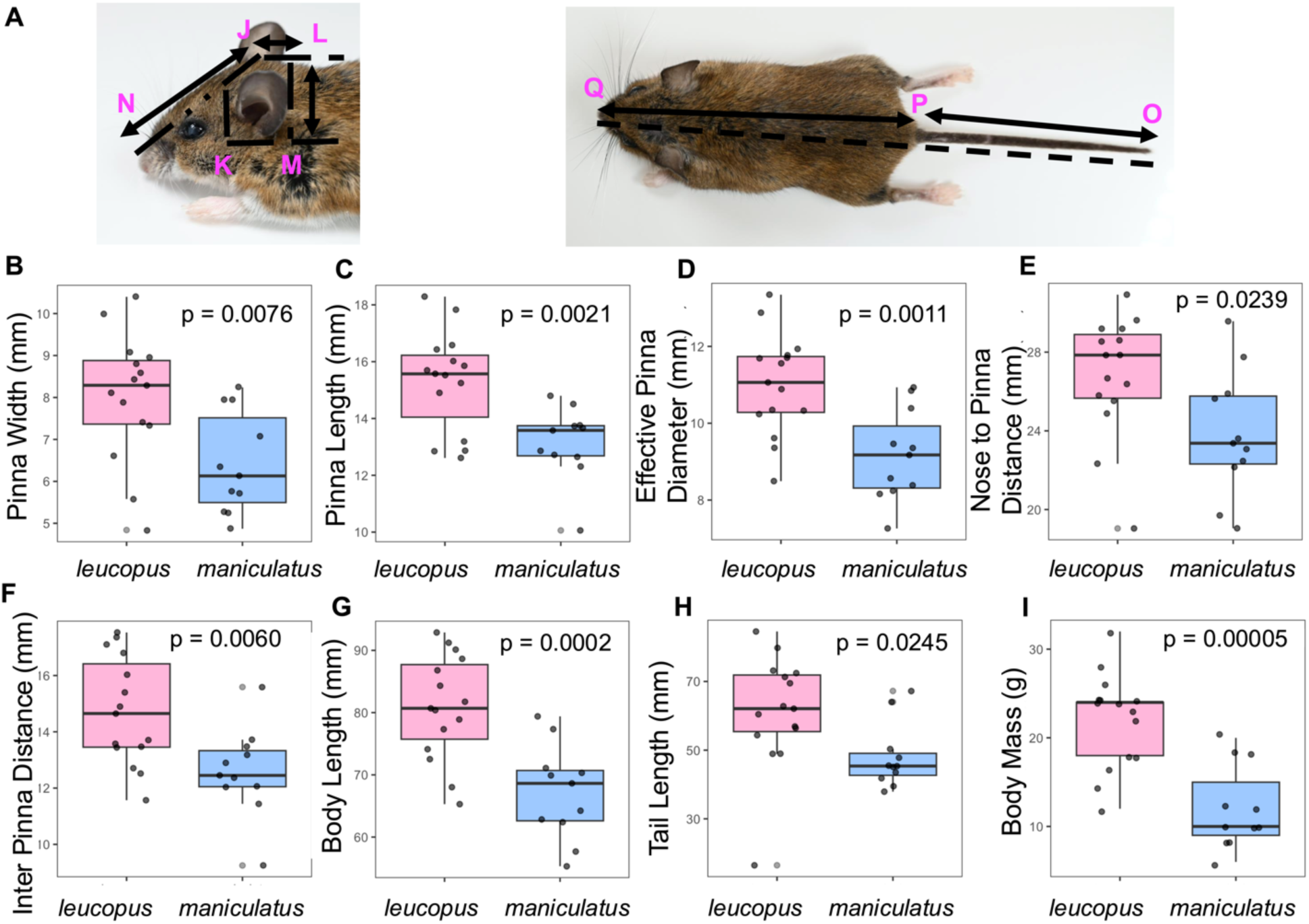
Morphological differences between *P. leucopus* and *P. maniculatus*. Pinnae, head, and body features (**A**) were evaluated between species (pink boxplot = *leucopus*, blue boxplot = *maniculatus*). Measurements JK show the inter pinnae distance, JN the nose to pinna distance, MK the pinna width, LM the pinna height, OP the tail length, and PQ the body length. Effective pinna diameter was calculated by taking the square root of pinna height multiplied by pinna width (MK/LM). Significant differences were observed for all features including Pinna width (**B**), Pinna length (**C**), Effective diameter (**D**), Nose to pinna distance (**E**), Inter pinna distance (**F**), Body length (**G**), Tail length (**H**), and Body mass (**I**). Peromyscus pictured is a wild caught *P. leucopus* captured in Payne County, Stillwater, Oklahoma. Image is presented only for demonstration of measurements.

### ABRs recordings

We recorded ABRs from wild *P. leucopus* and *P. maniculatus* using similar procedures as previous publications (Chawla and McCullagh, 2022; McCullagh et al., 2020; New et al., 2024). Rodents were sedated with an intraperitoneal injection of 60 mg/kg ketamine and 10 mg/kg xylazine for initial anesthesia followed by maintenance dosage of 25 mg/kg ketamine and 12 mg/kg xylazine. After being fully sedated, as indicated by lack of toe pinch reflex, the animals were transported to a small sound-attenuating chamber (Noise Barriers, Lake Forest, IL, USA), and positioned on a water pump heating pad (set to 37° C). Subdermal needle electrodes were inserted under the skin at midline between the ears over the brainstem (apex, active electrode), directly behind the apex on the nape (reference), and in the back leg of the sedated animals (ground electrode) for differential recordings. To obtain and amplify evoked potentials from electrodes positioned below the skin of the animal, we used a Tucker-Davis Technologies (TDT, Alachua, FL, USA) RA4LI head stage, a RA16PA preamplifier, and a Multi I/O processor RZ5 attached to a PC with custom Python software to record the data. Data were processed using a second order 50-3000 Hz filter and averaged across 10-12 ms of recording time over 500-1000 repetitions. Acoustic stimuli were delivered using TDT Electrostatic Speakers (TDT EC-1) for frequencies between 32 and 64 kHz or a TDT Electrostatic Speaker-Coupler Model (TDT MF-1) for clicks and frequencies ranging from 1 to 24 kHz. Speakers were coupled with custom ear bars fitted with Etymotic ER-7C probe microphones (Etymotic Research Inc., Elk Grove Village, IL) for in-ear calibration prior to recording for each animal. Tone stimuli were 4 ms in total duration with a 1 ms on-ramp and 1 ms off-ramp (2 ms ± 1 ms on/off ramps). Click stimuli were 0.1 ms in duration with alternating polarity. Acoustic stimuli were presented to the animal with a 30 ms interstimulus interval with a standard deviation of 5 ms (Laumen et al., 2016a). Randomizing interstimulus interval between each presentation has been shown to optimize the ABR waveform (Wang et al., 2020). Stimuli were produced at a sampling rate of 97656.25 Hz through a TDT RP2.1 real-time processor controlled by a custom Python program.

### ABR response threshold

ABR response thresholds were determined by using the visual technique outlined by (Brittan-Powell and Dooling, 2004). In short, threshold was defined to be between the intensity at which the waveforms were no longer present and the previous intensity at which they were visible in 5- and 10-dB increments specifically as absence of waveforms evoked by both left and right monaural stimuli (i.e. complete absence of both waves, 5 dB increments were used when near threshold). This method was used for analyzing the audiogram across frequencies presented (1, 2, 4, 8, 16, 24, 32, 46, 64 kHz) and intensities (90 - 10 dB SPL) in addition to click threshold.

### Monaural ABR recordings

Evoked potentials were recorded independently for each ear in response to single-ear click stimulation and measured based on peak amplitude (voltage from peak to absolute trough) and latency (time to peak amplitude) across the four peaks of the auditory brainstem recording waveforms (Chawla and McCullagh 2022). To calculate the monaural latency and amplitude for each species, we calculated the average of the monaural peak amplitude or latency of waves I and IV from the ABRs data obtained for sound presentation in each ear across intensities (60-90 dB SPL) (New et al., 2024; Zhou et al., 2006).

### Binaural auditory brainstem response recordings

To produce the binaural ABR response, we played broadband alternating polarity click stimuli simultaneously or with an ITD at 90 dB SPL to both pinnae of the sedated animal. The binaural interaction component (BIC) of the ABR was determined by subtracting the sum of the two monaural auditory brainstem evoked responses from the binaural ABR recordings (Benichoux et al., 2018; Laumen et al., 2016a). Custom Python software was used to measure the BIC amplitude and latency, with amplitude calculated to the baseline of the recording (Chawla and McCullagh, 2022). BIC was defined as the negative peak wave (DN1) at wave IV of the ABR after subtraction of the summed monaural and binaural responses (See New *et al*. 2024 for more details). To calculate how BIC varies with ITD, both species were presented with click stimuli with shifting ITDs of -2.0 to 2.0 ms in 0.5 ms steps. We calculated the peak latency and amplitude of DN1 at each ITD for each species. The ITD latency shift of the DN1 component of the BIC was determined in relation to the latency of DN1 at 0 ITD. The DN1 amplitude is highest at 0 ITD therefore amplitude for ITD shifts was transformed to relative amplitude with respect to 0 ms ITD to normalize recorded data (Laumen et al., 2016a). The average latency shift and relative DN1 amplitude values were used to make comparisons of binaural ABR as a function of ITD between species.

### Statistical analyses

All analyses and figures were created in R Studio version 4.0.3 (R Core Team 2020), using ggplot2 (Wickham, 2016) and lme4 (Bates et al., 2014) packages. Two-way analysis of variance (ANOVA) was used to statistically compare morphological characteristics between species. Log-transformed morphological features (pinna width, length, etc.) were compared with log body length and slope of the linear fit to describe potential allometry (slope > 1 indicating positive allometry and < 1 indicating negative allometry). Data were analyzed using linear mixed-effects models (LMMs) to account for repeat observation within one animal with species, sex, frequency, peak, percentage relative DN1 amplitude, and shifts in DN1 latency of the BIC as fixed effects and animal as random effect. If there were significant main effects of a variable of interest, estimated marginal means were used for pairwise comparisons (Russell, 2018). To control for multiple comparisons, emmeans implements a Tukey method for contrasts.

## RESULTS

Based on the NCBI GenBank species identification systems for COI, of the tail samples from the 26 animals collected, 15 individuals were identified as *P. leucopus*, and 11 as *P. maniculatus* (supplemental table 1, Figure 1). *P. maniculatus* body length ranged from 55 to 78 millimeters (mm) with mean body mass of 12 grams (g), while *P. leucopus* body length ranged from 65 to 93 mm with mean body mass of 21 g (Table 1).

### Morphological characteristics

Previous studies have shown that *P. leucopus* and *P. maniculatus* have significant differences in pinna sizes and craniofacial features (Choate, 1973; Light et al., 2021; Millien et al., 2017). We observed significant statistical differences for pinna attributes including pinna width (Df = 1, 24; F = 8.47; p = 0.0076) and pinna length (Df = 1, 24; F = 11.79; p = 0.0021) (Figure 2B and 2C respectively). In general, *P. leucopus* had longer and wider pinnae compared to *P. maniculatus*, with mean pinna length and width of 15.30 and 8.02 mm for *P. leucopus* and 13.15 and 6.42 mm for *P. maniculatus*. Similarly, effective pinna diameter (Df = 1, 24; F = 13.69; p = 0.0011) was significantly different between species, with *P. leucopus* having a wider effective pinna diameter compared to *P. maniculatus* (Table 2, Figure 2D). Craniofacial features including inter-pinna distance (Figure 2F; Df = 1, 24; F = 9.08; p = 0.0060) and distance from the nose to the pinna (Figure 2E; Df = 1, 24; F = 5.82; p = 0.0239) were significantly different between species with *P. leucopus* exhibiting a wider distance between pinnae and a longer distance from the nose to the pinna. Like pinnae morphology and craniofacial features, there were significant differences in body mass (Df = 1, 24; F = 24.2; p = 0.00005, Figure 2I), tail length (Df = 1, 24; F = 25.76, p = 0.0245, Figure 2H), and body length (Df = 1, 24; F = 18.32; p = 0.0002, Figure 2G) between both species, with *P. leucopus* weighing significantly more including longer tails and longer body length than *P. maniculatus* (Table 2). We tested if there were sex differences in craniofacial and pinna sizes in *P. leucopus*. There were no significant differences in craniofacial and pinna sizes between male and female *P. leucopus* (all p-value > 0.05). Sex differences were not explored for *P. maniculatus* due to limited number of female subjects of this species (9 males, 2 females). When anatomical data were compared for potential effects of body size (allometry), log features (pinna width, etc.) compared to log body length did not show positive allometry except for tail length, which indicated positive allometry (slope of 1.437 (*maniculatus*) and 1.218 (*leucopus*), supplemental Figure 2).

**Table 2:** Morphological characteristics features of *P. maniculatus* and *P. leucopus* of the Packsaddle wildlife management area (WMA), James Collin wildlife management area (WMA) and Payne County. Values presented represent the mean of different morphological features recorded ± standard error, the degrees of freedom, F-statistic, and p-value of morphological differences between species.

| **Morphological characteristics** | ***Maniculatus***<br>**mean ± S. E** | ***Leucopus***<br>**mean ± S. E** | **degrees of**<br>**freedom** | **F-statistic** | **p-value** |
| --- | --- | --- | --- | --- | --- |
| **Effective pinna diameter** | 9.16 ± 0.36 | 11.01 ± 0.34 | 1, 24 | 13.69 | 0.0011 |
| **Pinna length** | 13.15 ± 0.39 | 15.30 ± 0.45 | 1, 24 | 11.79 | 0.0021 |
| **Pinna width** | 6.42 ± 0.36 | 8.02 ± 0.39 | 1, 24 | 8.47 | 0.0076 |
| **Inter pinna distance** | 12.59 ± 0.47 | 14.72 ± 0.49 | 1, 24 | 9.08 | 0.0060 |
| **Nose to pinna distance** | 23.84 ± 0.96 | 26.82 ± 0.79 | 1, 24 | 5.82 | 0.0239 |
| **Body length** | 67.19 ± 2.28 | 80.87 ± 2.17 | 1, 24 | 18.32 | 0.0002 |
| **Tail length** | 48.01 ± 2.84 | 61.21 ± 4.21 | 1, 24 | 25.76 | 0.0245 |
| **Body mass** | 12.0 ± 1.0 | 22.0 ± 1.0 | 1, 24 | 24.2 | 0.00005 |

### ABR thresholds between species

Both *P. maniculatus* and *P. leucopus* displayed the best sensitivity to tones between 8 to 46 kHz, as indicated by lower ABR thresholds (below 55 dB SPL, Figure 3C). We detected no significant statistical difference in ABR frequency threshold between species across the frequencies tested (LMM, p = 0.543). Similarly, no significant difference in ABR frequency hearing threshold was observed between male and female *P. leucopus* (LMM, p = 0.523). We next investigated whether craniofacial or pinna measurements features are correlated with or influence ABR frequency thresholds in both species. We found that none of the morphological measurements had a significant effect on ABR frequency hearing thresholds between species (all p > 0.05) (supplemental materials, Figure 3,4,5).

**Figure 3:**
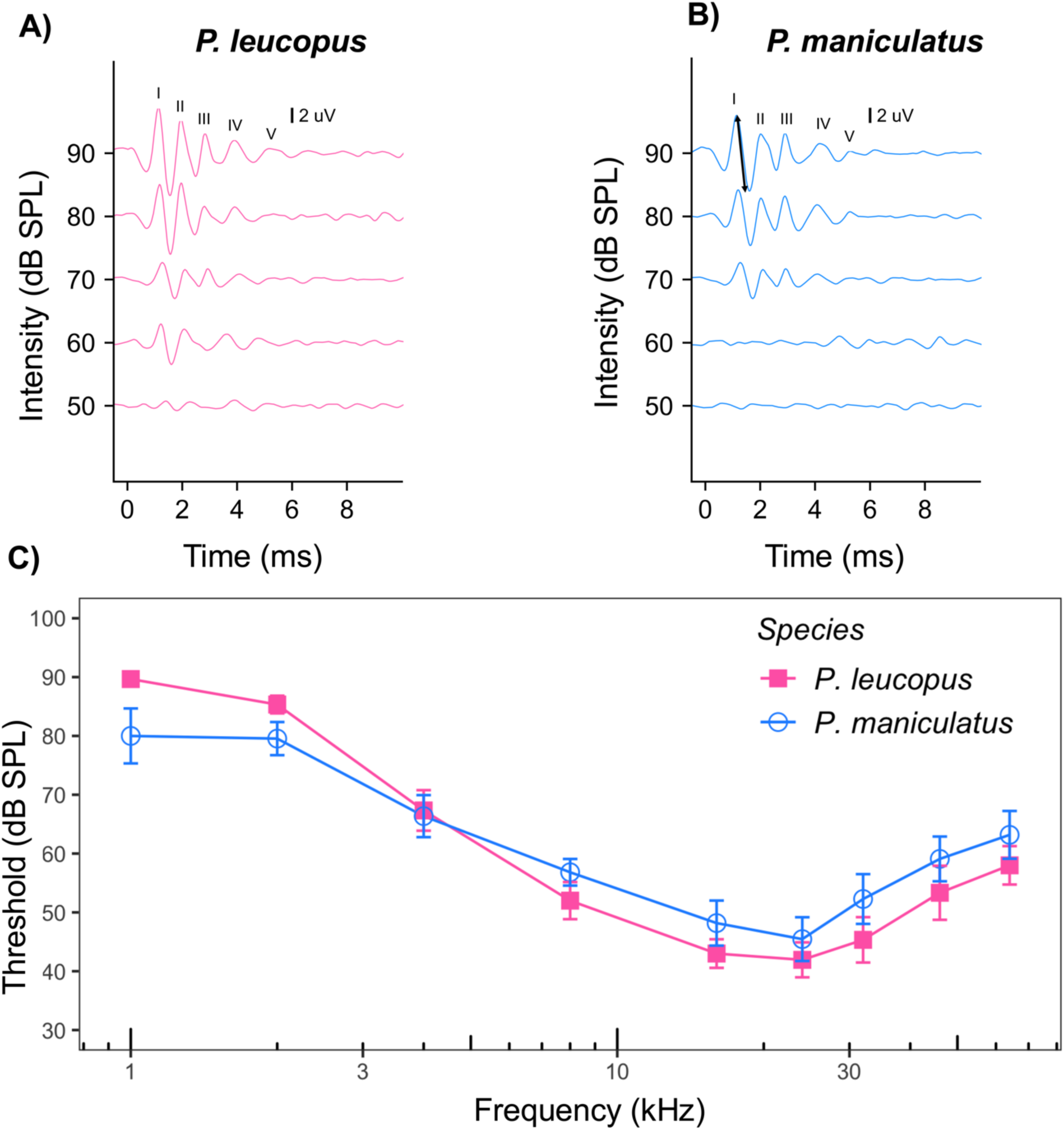
Figure 3A and 3B show auditory brainstem response patterns of a female *P. leucopus* (left) and a female *P. maniculatus* (right) determined with clicks of different intensities. Main waves I-V are identified in the 90 dB SPL example. Hearing range was measured across frequency (1-64 kHz) for both *P. leucopus* and *P. maniculatus* (Figure 3C). The vertical bars represent the standard error at each frequency. (Figure 3C). No significant main effects of frequency between species were found. Unfilled blue circles represent *P. maniculatus* while filled pink squares represent *P. leucopus*. Arrow in figure 3B shown how ampliture were calculated.

**Figure 4:**
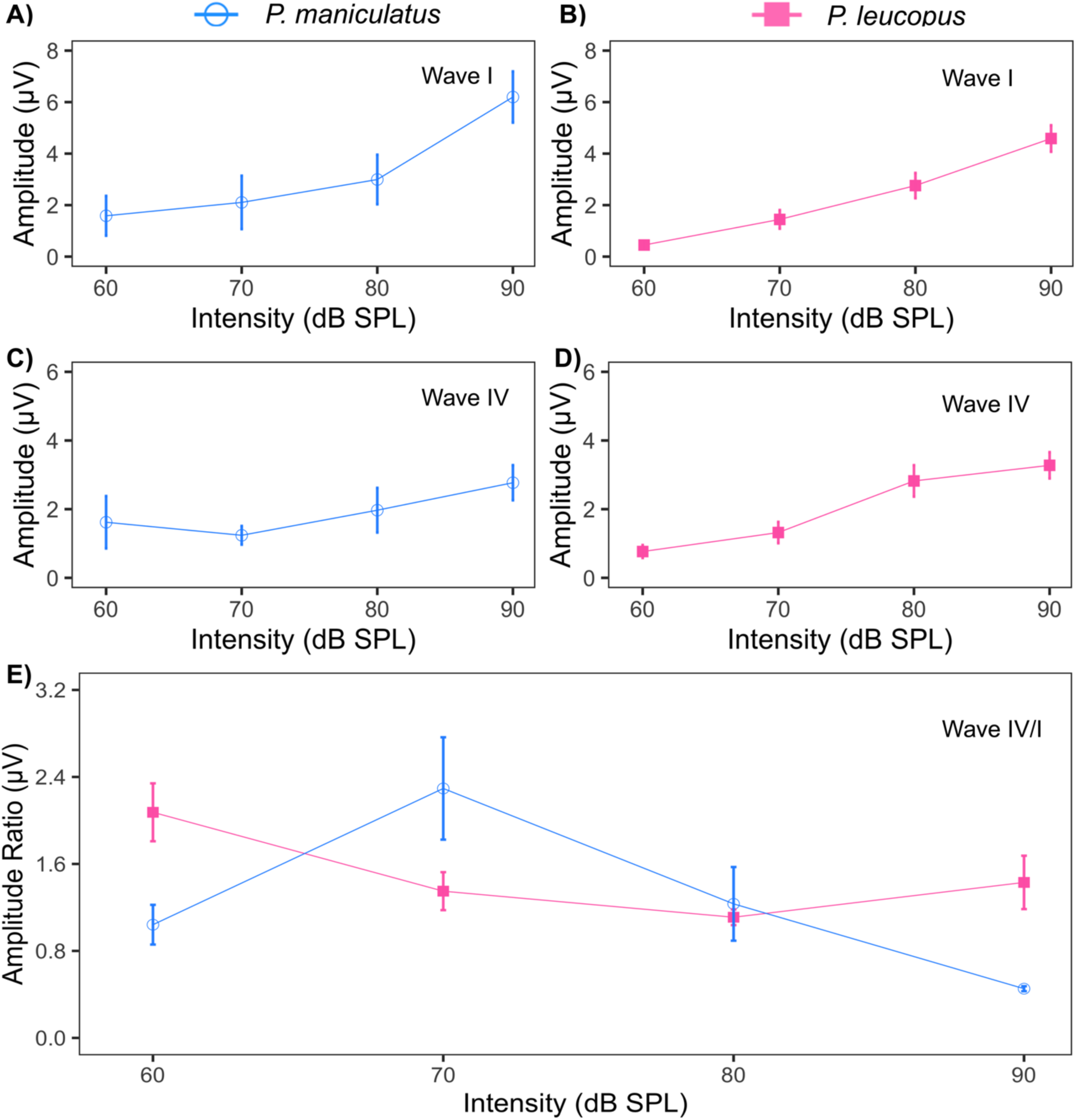
Average amplitude of wave I and wave IV of auditory brainstem responses determined with clicks of different intensities (pink filled square = *P. leucopus* (n = 15), blue unfilled circle = *P. maniculatus* (n = 11)). The average wave IV/I amplitude ratio at each intensity is shown in figure 4E. The vertical bars represent the standard error at each point. Significant main effects of intensity on wave I and IV amplitude were detected for both species.

### ABR waveform amplitudes

The amplitude of individual ABR waves was measured as the difference between the positive peak and the following negative trough (as shown by the arrow in Figure 3B). At 90 dB SPL, the average amplitudes of wave I and wave IV were 4.75 and 2.25 μV for *P. leucopus*. For *P. maniculatus*, the average amplitudes of wave I and wave IV were 5.87 and 2.13 μV at 90 dB SPL. We observed that the amplitude of waves I and IV (*P. maniculatus*: Figure 4A and 4C, wave I and IV unfilled circles; *P. leucopus*: Figure 4B and 4D, wave I and IV filled squares), increased monotonically with increasing intensity (60 to 90 dB SPL). Linear mixed-effect models revealed a significant main effect of intensity on the amplitude for both waves (LMM: p < 0.0001). However, no significant main effect of species was detected on the amplitude for both waves (LMM: wave I p = 0.289, wave IV p = 0.674). Similarly, a significant main effect of increasing intensity on peak amplitude for wave I and wave IV between male and female *P. leucopus* (LMM: p < 0.0001). However, there was no significant main effect of sex on monaural wave I and IV amplitude with increasing intensity for *P. leucopus* (LMM: p = 0.851).

### Monaural Amplitude Ratio

Monaural amplitude ratio was calculated by dividing the amplitude value of wave IV by the amplitude value of wave I for left and right pinnae at each intensity. For both species, wave IV (4C and 4D) overall had smaller amplitude than wave I (Figure 4A and 4B) and as a result, the wave IV/I amplitude ratio was generally lower than 1.0 at most intensities tested in both species (Figure 4E, *P. leucopus*: filled squares and *P. maniculatus*: unfilled circles). The results of the linear mix-effect model revealed no significant main effects of either intensity (LMM: p = 0.158) or species (LMM: p = 0.567) on the amplitude ratio of wave I and IV. Similarly there was no main effect of either intensity (LMM: p = 0.465) or sex (LMM: p = 0.705) for *P. leucopus* on the amplitude ratio of wave I and IV.

### Absolute Latency

Peak latency is the time interval between the presentation of a sound stimulus and the peak at maximum amplitude of the designated wave. We calculated the average peak latency of waves I and IV of both studied species across click intensities (60 to 90 dB SPL). Linear mixed-effect models revealed a significant decrease in peak latency for both waves with increasing intensity between species (Figures 5A, 5B, 5C and 5D, LMM: all p < 0.0001). There was a significant main effect detected for species in peak latency for wave I (LMM: p = 0.00276), but not wave IV (LMM: p = 0.0797) with increasing intensity. However, pairwise comparisons did not show significance at a particular intensity suggesting that overall *P. maniculatus* had shorter latencies, when considering all intensities, than *P.* leucopus for wave I only when using non-relative measures (Figure 5A, 5B). Similarly, significant decreases in peak latency with increasing intensity were observed for wave I and wave IV between male and female *P. leucopus* (LMM: all p < 0.0001). There was a significant main effect of sex detected for wave I in *P. leucopus* (LMM: p = 0.0106), driven by sex differences at 90 dB (LMM: p = 0.0027), with no main effect of sex on wave IV (LMM: p = 0.293). We next determined the slope latency of ABR waves between species by plotting the latency data of wave I and IV against different click intensities of each species (Supplemental, Figure 6). The slope of wave I latency was - 0.02 for *P. leucopus* and – 0.01 for *P. maniculatus*, while the slope of wave IV was – 0.03 for *P. leucopus* and – 0.04 for *P. maniculatus*.

**Figure 5:**
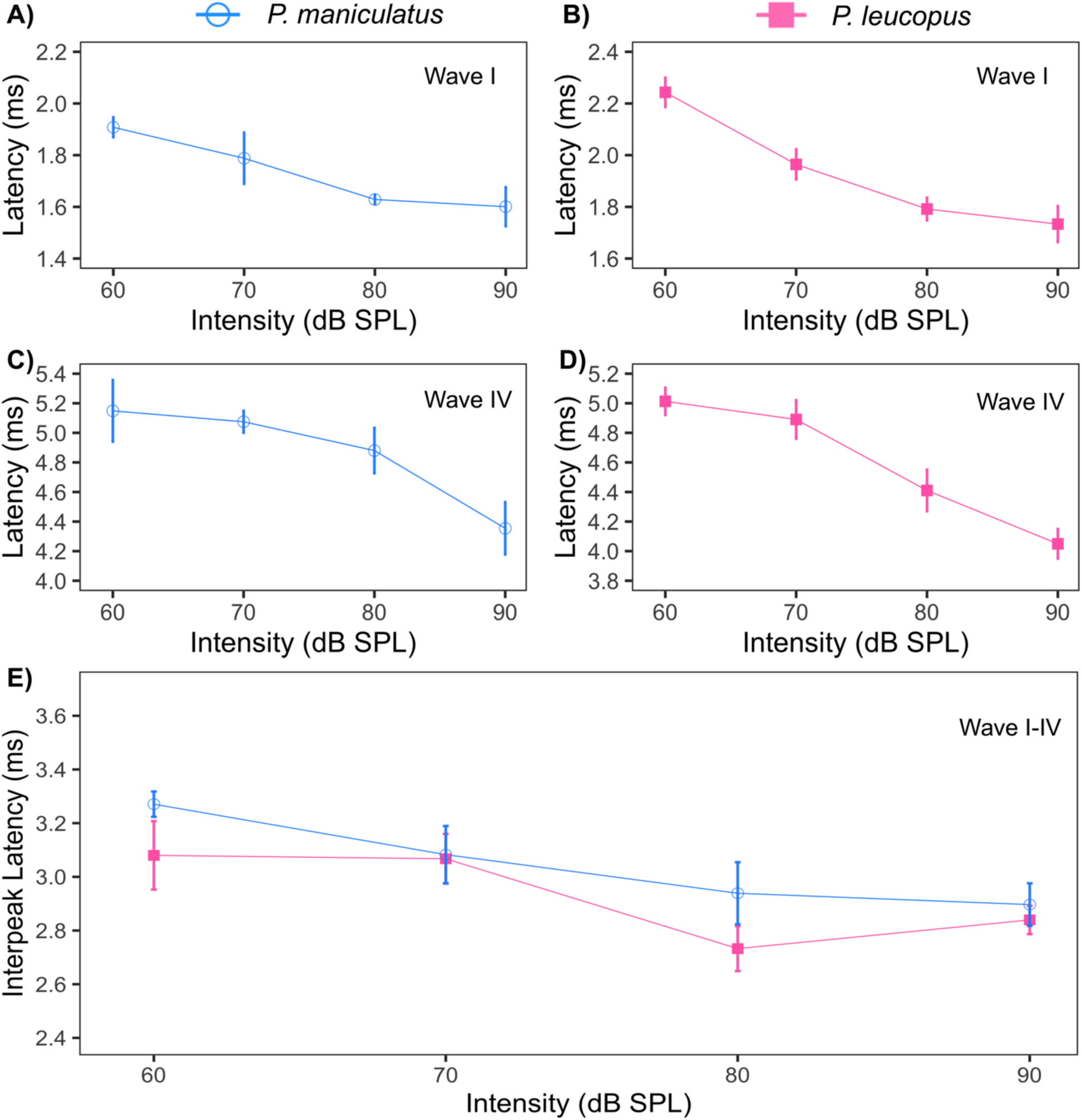
Average peak latency of wave I and wave IV of auditory brainstem responses determined with clicks of different intensities (Pink filled squares = *P. leucopus* (n = 15), Blue unfilled circles = *P. maniculatus* (n = 11)). The average wave I-IV inter-peak latency at each intensity is shown in 5E. The vertical bars represent the standard error at each point.

### Inter-peak latency

Inter-peak latency was calculated as the difference in latency from the wave I peak to wave IV peak for left and right pinnae at each intensity, which provides a more relative measure for comparing between species. Because the decrease in peak latency with increasing intensity was more pronounced for wave IV than for wave I, the inter-peak latency between wave I and wave IV also showed a significant decrease as intensity increased, as illustrated in Figure 5E (LMM: p = 0.00155, pink filled squares = *P. leucopus*, blue unfilled circles = *P. maniculatus*). However, no main effect of species was detected on the inter-peak latency of waves I and IV between both species (LMM: p = 0.281). There was a significant decrease in wave I-IV inter peak latency with increasing intensity for male and female *P. leucopus* (LMM: p = 0.0151) however there was no main effect of sex (LMM: p = 0.252).

### Binaural ABR measures

We used the latency shift of the DN1 component of the BIC and relative DN1 amplitude to show the relationship of ITD on latency and relative amplitude of the BIC across species. For both species, DN1 latency increased with longer ITDs and DN1 relative amplitude decreased with longer ITDs. The average DN1 amplitudes at 0 ITD were 2.72 µV and 1.74 µV for *P. maniculatus* and *P. leucopus*, respectively. The average latency for the DN1 component at 0 ITD was 5.0 ms for *P. maniculatus*, compared with 5.6 ms for *P. leucopus*. Linear mixed-effects models indicated significant main effects of ITD across relative DN1 amplitude (LMM: p < 0.0001), indicating that as ITD increased, relative DN1 amplitude decreased. No main effect of species was observed across relative DN1 amplitude (Figure 6B, LMM: p = 0.818). Similarly, there were significant main effects of ITD across relative DN1 amplitude between male and female *P. leucopus* (LMM: p < 0.0001), while no significant main effect of sex was detected (LMM: p = 0.0929). There were statistically significant main effects of both ITD (LMM: p < 0.0001) and species (LMM: p < 0.0162) in latency shift of the DN1 component of the BIC in relation to ITD normalized for 0 ITD between *P. maniculatus* and *P. leucopus* (Figure 6A). Shift in DN1 latency of the BIC was significantly faster in *P. maniculatus* compared to *P. leucopus* at 1.0 ms (p = 0.0377) and 2.0 ms (p = 0.0222) ITD (Figure 6A). There were significant differences in in latency shift of the DN1 component of the BIC in relation to ITD normalized for 0 ITD for male and female *P. leucopus* (LMM: p < 0.0001) but no main effect of sex (LMM: p = 0.843).

**Figure 6:**
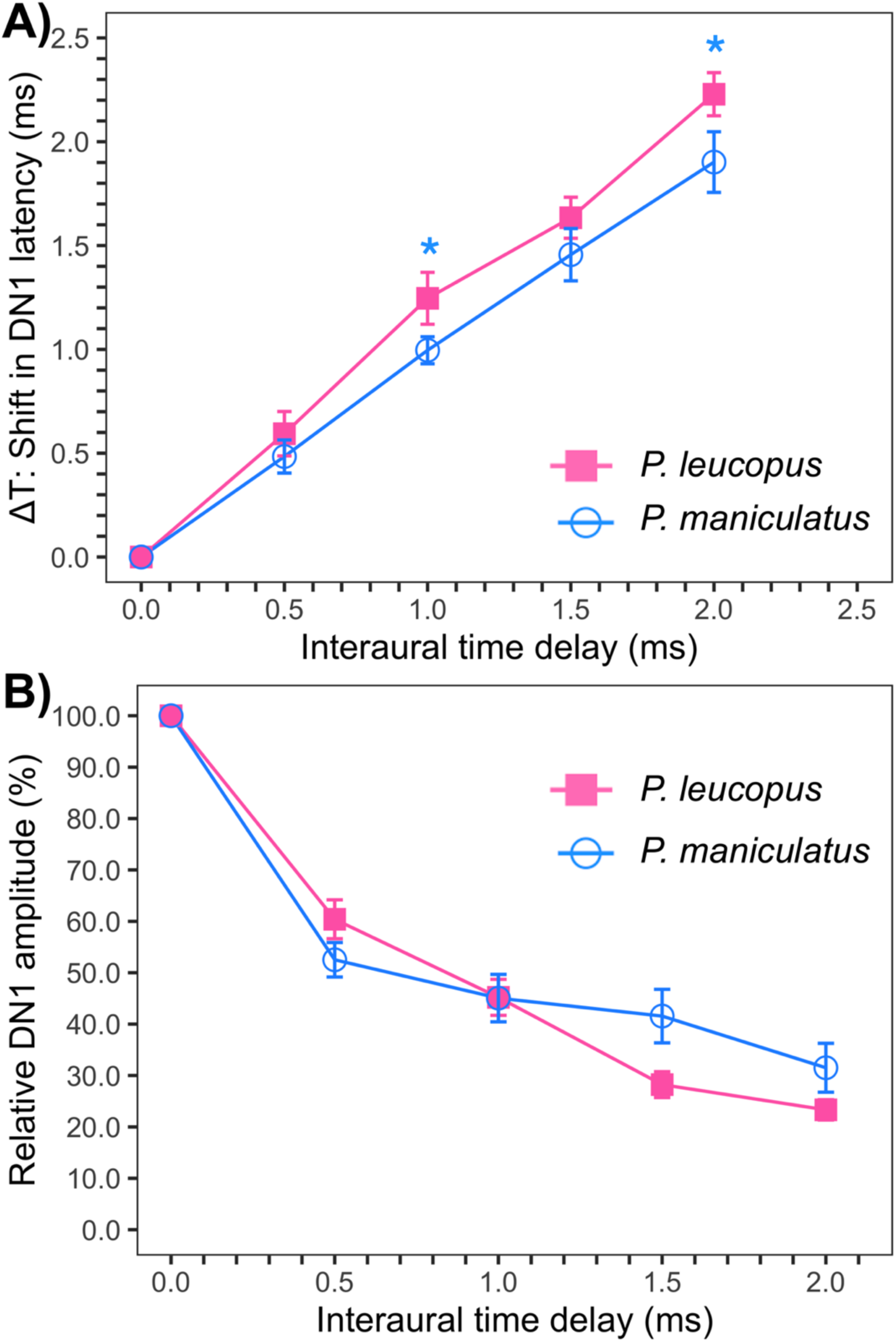
Binaural ABRs in wild *P. leucopus* (pink filled square) and *P. maniculatus* (blue unfilled circle). 6A, Shift in DN1 latency (ms) relative to ITD; reference latency at ITD = 0 is set to 0 ms. 6B, percentage relative DN1 amplitude relative to ITD normalized to the DN1 amplitude for ITD = 0 ms. Relative amplitude and latency of the DN1 BIC with varying ITD between - 2 to + 2 ms in 0.5 ms steps were measured. Significant main effects of ITD and species were detected in BIC shift in DN1 latencies between both species. No significant main effect of species was observed for relative amplitude of the BIC across ITDs. The vertical bars represent the standard error at each ITD.

## DISCUSSION

In this study, we used craniofacial morphology, pinna features, and ABRs to compare morphological features important for hearing with physiological measures of ABR amplitude and latency of two species of the genus *Peromyscus*. Like previous findings (Choate, 1973; Light et al., 2021; Millien et al., 2017), we detected significant morphological differences between species for all measurements (including pinna length, pinna width, etc.) with *P. leucopus* displaying larger features. ABR-derived detection thresholds revealed that both species share similar ABR response thresholds across frequencies with the lowest ABR frequency thresholds between 8-46 kHz, which is in agreement with previous findings that showed *Peromyscus* species have best hearing sensitivity between 8-16 kHz (Capshaw et al., 2022; Dice and Barto, 1952; Ralls, 1967). Significant main effects of intensity were detected in monaural amplitude of ABR wave I and IV between both studied species, which is similar to Zhou and colleague’s findings using laboratory mouse strains (Zhou et al., 2006). Measurements of the BIC indicated significant main effect of ITDs on DN1 amplitude and latency between the two species. Significant main effect of species was detected for DN1 latency, but not for DN1 amplitude. Overall, our results revealed that both species have similar ABR hearing thresholds with *P. maniculatus* having shorter latency BIC and smaller anatomical features compared to *P. leucopus*.

Morphological features including cranial size and shape, size of the pinnae, body and tail length differ widely within species of the genus *Peromyscus* (Light et al., 2021; Ordóñez-Garza et al., 2010). Our data showed differences in all measured anatomical traits between species consistent with previous studies (Choate, 1973; Light et al., 2021; Millien et al., 2017). Previous studies used two-dimensional (2D) geometric morphometrics (Light et al., 2021), and micro-CT (Riede et al., 2022) as tools to morphologically differentiate rodents of the genus *Peromyscus*.

Light *et al* (2021) showed differences between *P. leucopus* and *P. maniculatus* based on head size, pinnae features, hindfoot length, and other morphological features similar to our study. Consistency in morphological features (pinna length, pinna width, and body weight) documented in this study provide additional evidence supporting the use of these morphology traits as reliable indicators for distinguishing species within the genus *Peromyscus*. Using ABRs, Capshaw *et al*. (2022) observed low hearing sensitivity to frequencies below 2 kHz in two laboratory *Peromyscus* species (*P. leucopus* and *P. californicus*). Our findings are consistent with Capshaw *et al*. (2022), with hearing thresholds around 85 dB SPL at frequencies below 2 kHz in both studied species, suggesting relatively poor hearing sensitivity of both studied species to low frequencies. Small-headed mammals generally are not as sensitive to low frequencies and do not generate significant directional information with them due to minimal differences in timing and intensity between pinnae (Lauer et al., 2018). Therefore, it is thought that small mammals rely on high frequencies for directional hearing with exception of some subterranean mammals including the naked mole-rat (*Heterocephalus glaber*), the plain pocket gopher (*Geomys bursarius*), and the blind mole rat (*Spalax ehrenbergi*) that have poor sound localization and lack high frequency hearing (Heffner and Heffner, 1993, 1992, 1990), though see (Barker et al., 2021; Gessele et al., 2016; McCullagh et al., 2022). The limited ability of small mammals (with exception of Mongolian gerbils, chipmunks, groundhogs, hamsters, and others), like members of the genus *Peromyscus*, to detect low frequency sounds has been attributed to selective pressure linked with the absence of cues for localizing sounds in the horizontal plane for these small-headed animals (Heffner et al., 2001). Therefore, it is not surprising that we did not observe low frequency sensitivity for the two studied species in the current investigation.

*P. leucopus* and *P. maniculatus* are both highly territorial and produce both sonic and ultrasonic vocalizations between 0.8 to 28 kHz (sustained vocalizations: frequency ranges between 10-25 kHz, sweep vocalization: frequencies above 25 kHz, and barks: frequency ranges between 0.8 and 6 kHz) (Miller and Engstrom, 2012; Pomerantz and Clemens, 1981; Riede et al., 2022). The frequency ranges of ultrasonic vocalizations of both studied species correlate with their frequency thresholds (Figure 3C, best frequency threshold ranging from 8 - 46 kHz). Related species’ (California mouse, *P. californicus*) defensive and distress vocalizations are known to be associated with sounds ranging from 2 -30 kHz (Rieger and Marler, 2018). While limited studies have described distress and defensive vocalizations across the genus *Peromyscus*, previous investigations have reported that members of this genus produce agonistic calls such as chits and barks at frequencies between 6 to 15 kHz (Houseknecht, 1968; Pasch et al., 2017). These findings suggest a good match of *Peromyscus’s* vocalization with their ABR threshold sensitivity (8 – 46 kHz) likely contributing to vocal air-borne communication in the wild.

The white-footed mice (*P. leucopus*) and deer mice (*P. maniculatus*) occur throughout Oklahoma but generally occupy different habitats, with *P. maniculatus* being more common in grasslands and *P. leucopus* primarily inhabiting shaded forests (Hackney and Stancampiano, 2015; Stancampiano and Schnell, 2004). In our study, *P. leucopus* subjects were mainly captured in shrubland and forested habitats, while *P. maniculatus* subjects were found in open grassland habitats. Our findings revealed that *P. leucopus* has similar sensitivity to frequencies as *P. maniculatus* across all frequencies tested. We may be missing potential species level difference however due to suspected differences in age between the species. Weight distribution suggests that eight of the 11 *P. maniculatus* subjects in the current study were juveniles, while 10 of the 15 *P. leucopus* were adults. Age differences may mask any potential species level differences, as small shifts in audiogram thresholds have been observed in *P. leucopus* with aging (Capshaw et al., 2022). A comparative study evaluating the vocalization content and sound propogation of vocalizations of both species in their respective habitats, across different age groups, would shed novel insights into how habitat-related factors and age might influence sound reception and communication strategies both within and among closely related *Peromyscus* species.

Amplitude of wave I and IV tend to increase monotonically in most small mammals with increasing intensity when measured by click stimuli (Zhou et al., 2006). Similar patterns have been reported in other taxa commonly used in evoked potential studies (Backoff and Caspary, 1994; Neil J. Ingham, 1998). Though we didn’t see any difference between the species for relative wave amplitude, we did observe differences in craniofacial size and mass, however they were proportional (larger animals had more mass and bigger pinna). Previous studies indicate that smaller craniofacial size with small body mass may bring the recording electrodes into closer proximity to the generators, resulting in larger amplitudes compared to those with large body mass (Merzenich et al. 1983) though in our studies the disparities in size may not have been large enough for this effect to be seen.

ABR wave amplitude can be affected by several factors including electrode position, animal body temperature, external noise, recording protocol, and equipment characteristics, therefore normalization between waves can help control for this variability. In humans, it has been shown that auditory deficits related to retrocochlear pathology may lead to a decrease in wave IV amplitude, and ultimately cause a decrease in the wave IV/I amplitude ratio (Arnold 2000). Our data revealed that the wave IV has a smaller amplitude than wave I in both species at most intensities tested, resulting in a wave IV/I ratio less than 1.0 (Figure 4E). Previous work measuring ABR in inbred mouse strains, rats, gerbils, cats, guinea pigs, and humans indicated that the ABR waves I and II are generally larger amplitude than ABR waves III and IV, which is in agreement to the current study (Moore, 1983). However, wave II and III are relatively larger in rats and guinea pigs, while shifted to wave IV in cats and wave IV-V complex in humans (Merzenich et al., 1983). Accordingly, the species-specific differences in individual ABR wave amplitude may result from complex factors including the evolution of the central nervous system, neuronal response characteristics within the brainstem, and neural conduction velocity.

ITD and ILD are two cues that animals with external pinnae use for sound localization. ITDs are generally processed by neurons in the medial superior olive (MSO <2kHz) while ILDs are mainly processed by neurons in the lateral superior olive (LSO >2 kHz) (Grothe et al., 2010; Suzuki and Horiuchi, 1981). Previous studies reported that the mean DN1 amplitude at 0 ITD was 0.2 µV in humans, about 5 µV in guinea pig, 1.8 µV in gerbil, and 2.3 µV in cats (Goksoy et al., 2005; Jones and Van der Poel, 1990; Laumen et al., 2016a; Riedel and Kollmeier, 2006; Ungan et al., 1997). Comparatively, the DN1 amplitude at 0 ITD of wild *Peromyscus* species is similar to that of gerbils and cats, higher than that of humans and significantly lower than that of guinea pig. Differences in DN1 amplitude at 0 ITD observed could be a result of smaller distance of the recorded electrodes to the brainstem in small animals, as well as electrode configuration, or other procedural differences.

Numerous publications have reported that latency of the DN1 component in humans ranges from 5.6 to 6.8 ms, while those of other animal models (gerbil, cats, guinea pig) range from 3.7 to 4.8 ms (Goksoy et al., 2005; Jones and Van der Poel, 1990; Riedel and Kollmeier, 2006; Ungan et al., 1997). Our data on the latency DN1 component is consistent with latencies observed in humans and is somewhat slower than what is seen in other animal models. Our results are similar with others that show the latency of the DN1 component increases with longer ITDs in cats, gerbil, guinea pig, and humans (Goksoy et al., 2005; Laumen et al., 2016b; Riedel and Kollmeier, 2006; Ungan et al., 1997). Indeed, it has been suggested that the increase in DN1 latency with increasing ITD reflects the anatomy and interaction between excitatory and inhibitory neurons in the superior olivary complex (Karino et al., 2011).

We observed faster latencies of DN1 and wave I in *P. maniculatus* compared to *P. leucopus*, (when not considering the relative interpeak latency for wave I). It is hard to speculate whether the difference in DN1 latency observed between both species is associated with head size or the number of cells in the superior olivary complex (SOC) nuclei. Studies characterizing the number of excitatory and inhibitory cells in the SOC of both species would be beneficial to allow for evaluation of the effects of head size or medial superior olive (MSO) and lateral superior olive (LSO) size in shifts of the DN1 latency among *Peromyscus* species. Further studies involving more *Peromyscus* species and other techniques, such as head-related transfer functions, are needed to assess if larger external pinna sizes contribute to additional features of *Peromyscus* hearing such as the use of spectral notches and the contribution of the pinna to horizontal cues like ITD and ILD, particularly since our in-ear presentation of ITD stimuli bypass the pinna. We calculated the functional interaural distance for each species by summing the mean inter-pinna distance and pinna width divided by the speed of sound in air to evaluate the availability of ITD cues for each species. While this technique is limited due to our use of calipers and is not exactly the same as the time delay caused by sound traveling around the head, we nonetheless used this to roughly estimate the functional interaural distance for each species. We found that *P. maniculatus* have a shorter functional interaural distance (± 55 μs) compared to *P. leucopus* (± 67 μs) which is consistent with smaller heads in *P. maniculatus*.

There are some limitations to the techniques employed in this study. Calipers are less accurate as features get smaller due to their measurement sensitivity, therefore measures of pinna and head morphology are likely to be less accurate than larger measurements such as body length and tail length. We conducted analyses correcting for overall body length; however, they did not show significant positive allometry (slope > 1) indicating that either these features were not allometric or the loss of accuracy of measurements at smaller distances contributed significantly to error. However, the one measurement that showed positive allometry was tail length, which is one of the longer, or perhaps more accurate, measures suggesting that finer measurement tools might be needed to make further arguments about effects of overall body size and morphological features on hearing in these species. There are also limitations to using ABRs as measures of hearing, including that interpretation of thresholds using visual observation, as performed in our study, can be subjective (Suthakar and Liberman, 2019). However, others have shown minimal differences between algorithms and observers to auditory threshold measurements (Capshaw et al., 2022). Further validation of our observer method with more quantitative algorithms would be useful to confirm threshold values reported here, though our thresholds coincide well with the published literature in one of these species (Capshaw et al., 2022). Lastly, behavioral measures of hearing can show differences compared to ABRs, and indeed anesthetics used, montage of electrodes, calibration of sounds (in ear or other methods), sound presentation, and other factors all may influence ABR results making cross-species and cross-publication results difficult to interpret (Ramsier and Dominy, 2010; Wolski et al., 2003). However, the current study used the same parameters across both species and showed results consistent with the literature and what might be expected for species that are closely related but differ primarily in size giving us confidence in the results presented here.

## CONCLUSIONS

Our findings provide a deeper understanding of auditory similarities and differences between two species of *Peromyscus* and validate that the highly abundant *Peromyscus* may serve as a future model for auditory studies. Both species show differences in craniofacial and pinna features and exhibit best ABR thresholds at frequencies ranging from 8 to 46 kHz. *P. maniculatus* showed shorter relative latencies of the DN1 component of the BIC, while relative DN1 amplitude was not different between the species. Further physiological assessment exploring hearing between the sexes at different ages and across the lifespan are needed to further show whether there are differences in hearing under these conditions. In addition, clarifying the role of the BIC between sexes across species of the genus *Peromyscus* is important to understand its relevance for potential sex differences.

## ACKNOWLEDGMENTS

We would like to thank Game wardens, Benny Farrar and Marcus Thibodeau for housing and giving us opportunities to sample at James Collin and Packsaddle Wildlife Management Areas. Also, we would like to thank members of team wild rodent of the McCullagh lab which helped in trapping and performed ABRs. We would also like to thank Dr. Tim Lei and Benzheng Li for their creation of the ABR acquisition and analysis custom python software. Dr. Fabio Machado helped with interpreting analyses of allometry and body size measurements. NSF RaMP DEB 2216648, and Oklahoma State University College of Arts & Sciences (CAS) Research Program to EAM helped fund summer support, RA support for LJ and undergraduates involved in the project, and some materials (CAS research award). Support for VYF was provided by a Wentz and CAS AURCA program support, and EMN with Wentz fellowship support as well as additional support for LJ from the Payne County Audubon Society.

## DATA AVAILABILITY

The data of the study will be made available upon request.

## AUTHOR CONTRIBUTIONS

LJ, EMN, TCW, VYF, And DMJ captured the animals and LJ and EMN collected the ABR data for the manuscript. LJ, DMJ, and GOUW performed DNA analysis on tails snip samples while BL helped with data analysis and interpretation. LJ and EAM completed the statistical analysis and developed the idea of the paper. LJ wrote the manuscript and all other authors revised and edited the manuscript.

## COMPETING INTERESTS

The authors declare no competing interests.

## SUPPLEMENTARY MATERIAL

See supplementary materal at [] for access to supplementary figures and tables.

## Notes

### Competing Interest Statement

The authors have declared no competing interest.

### Summary of Updates

We have made relative measures of auditory brainstem monaural and binaural components.

