## supplemental figures and tables for "Auditory Brainstem Responses in Two Closely Related *Peromyscus* Species (*Peromyscus leucopus* and *Peromyscus maniculatus*)"

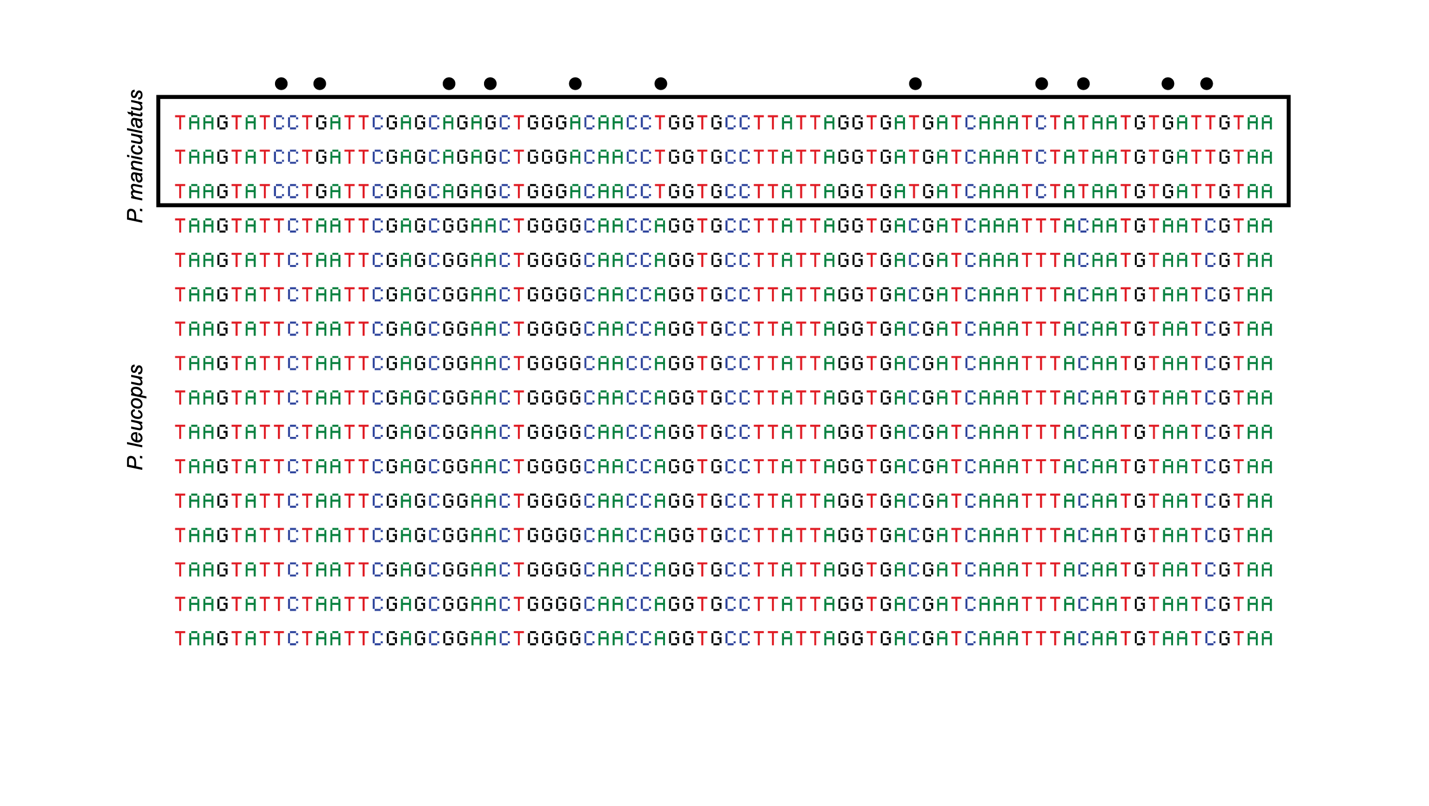

**Figure 1:** Fragment of the alignment of three random *P. maniculatus* sequencies (top box) aligned with 13 randomly selected *P. leucopus* (bottom) samples. Base pair changes that help indicate species differences are highlighted with black dots at the top.

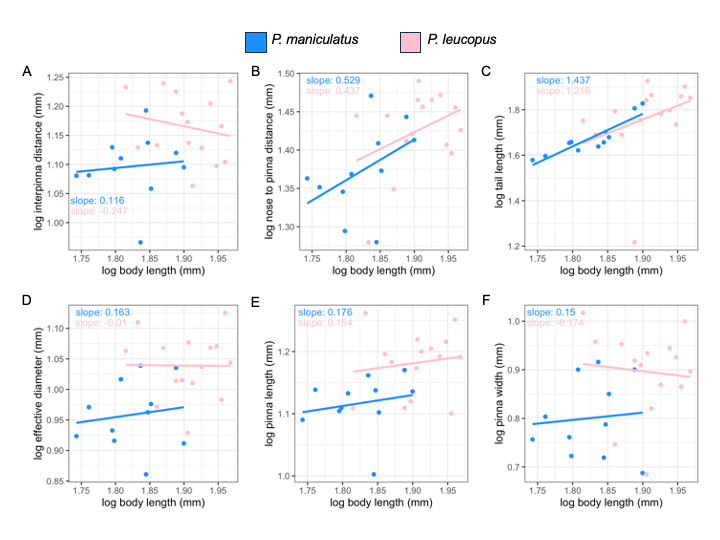

**Figure 2:** log values of traits

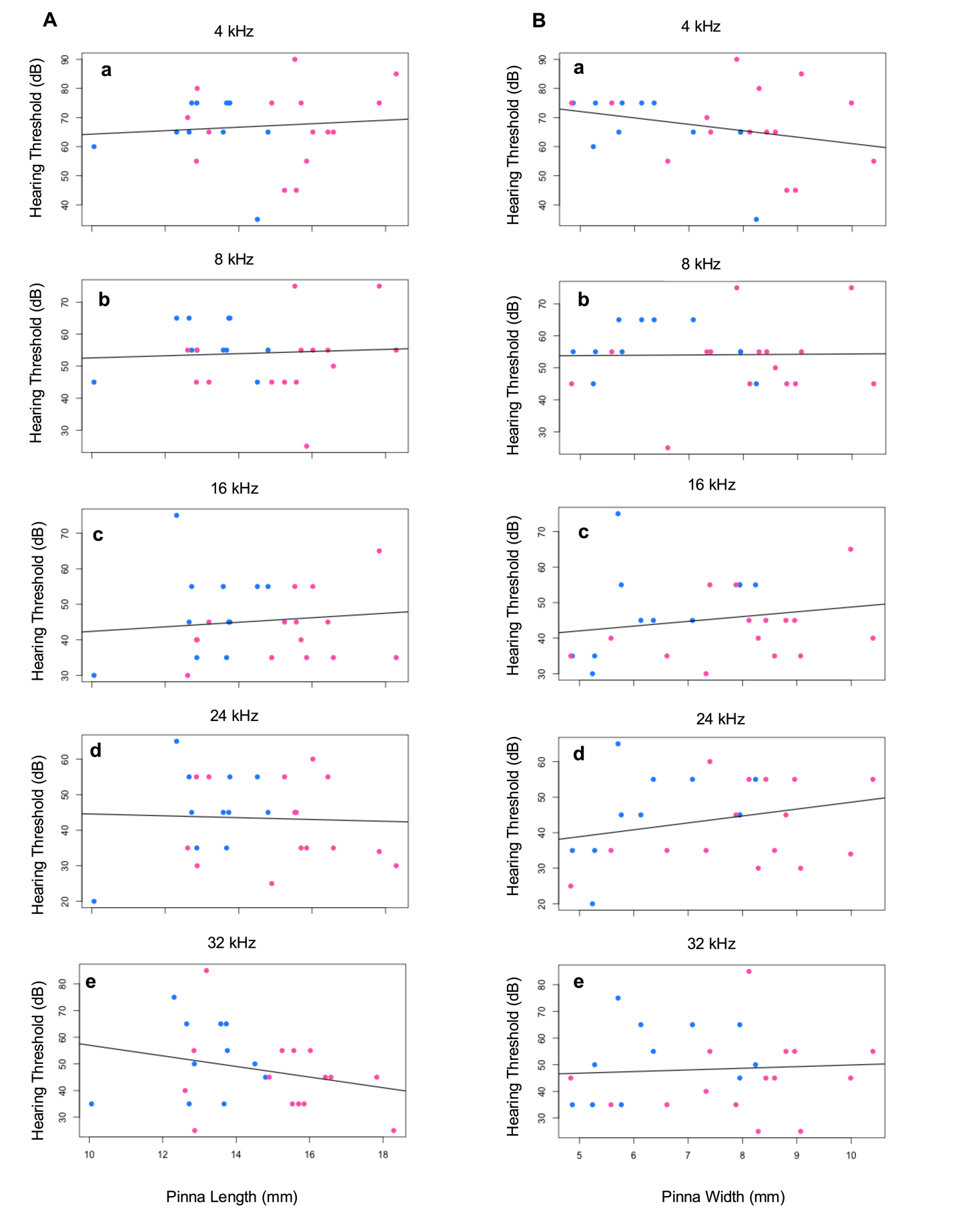

**Figure 3:** Scatter plots of frequency thresholds and morphological features (A = Pinna Length, B = Pinna Width) between species. Blue dots represent *P.* *maniculatus* and pink dots represent *P. leucopus*. No significant effects of pinna length and pinna Width were detected on best frequency hearing thresholds between species at 4, 8, 16, 24, and 32 kHz (All Pvalue < 0.05)

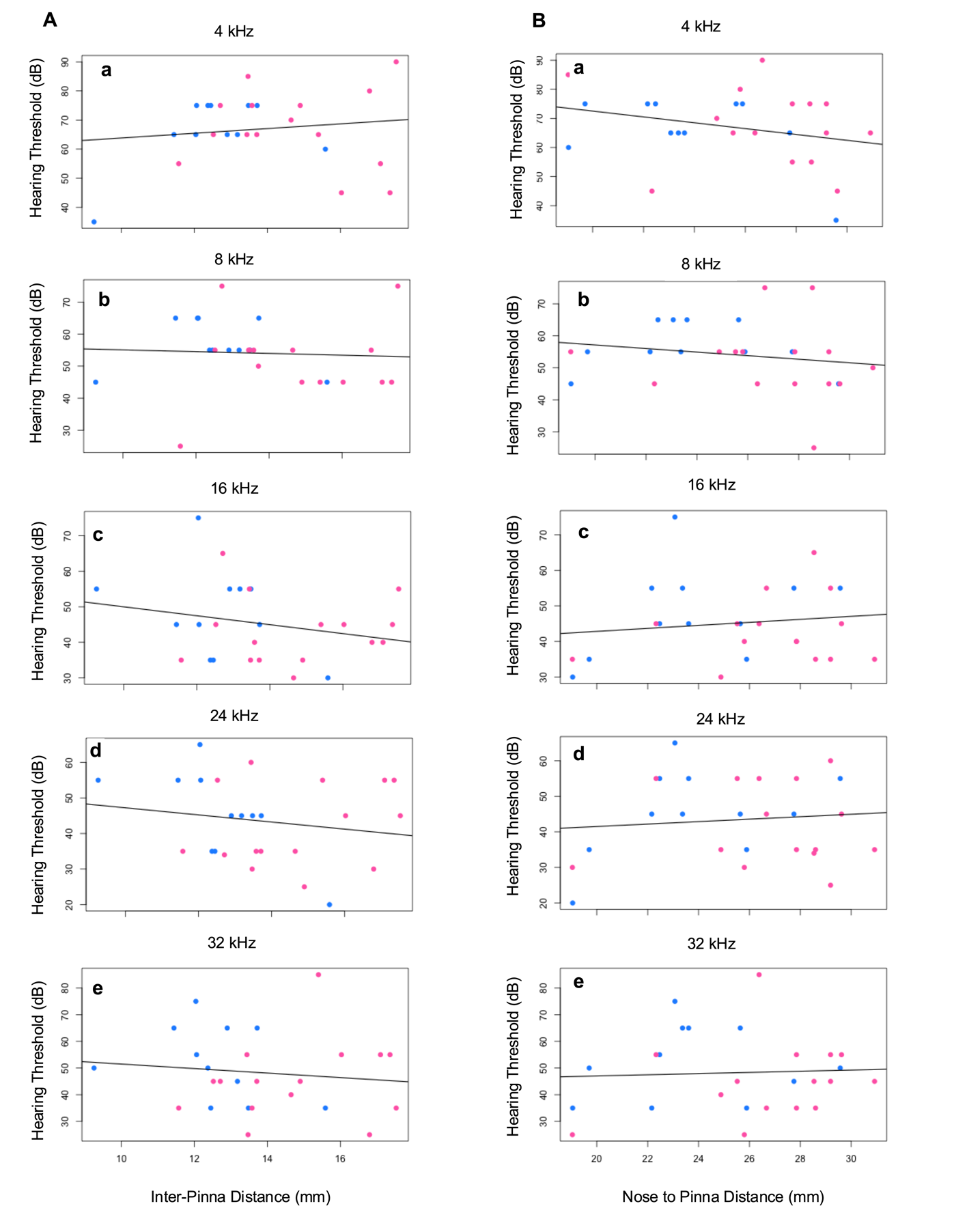
**Figure 4:** Scatter plots of frequency thresholds and morphological features (A = Inter-Pinna Distance, B = Nose to Pinna Distance) between species. Blue dots represent *P.* *maniculatus* and pink dots represent *P. leucopus*. No significant effects of inter-pinna distance and Nose to pinna distance were detected on best frequency hearing thresholds between species at 4, 8, 16, 24, and 32 kHz (All Pvalue < 0.05)

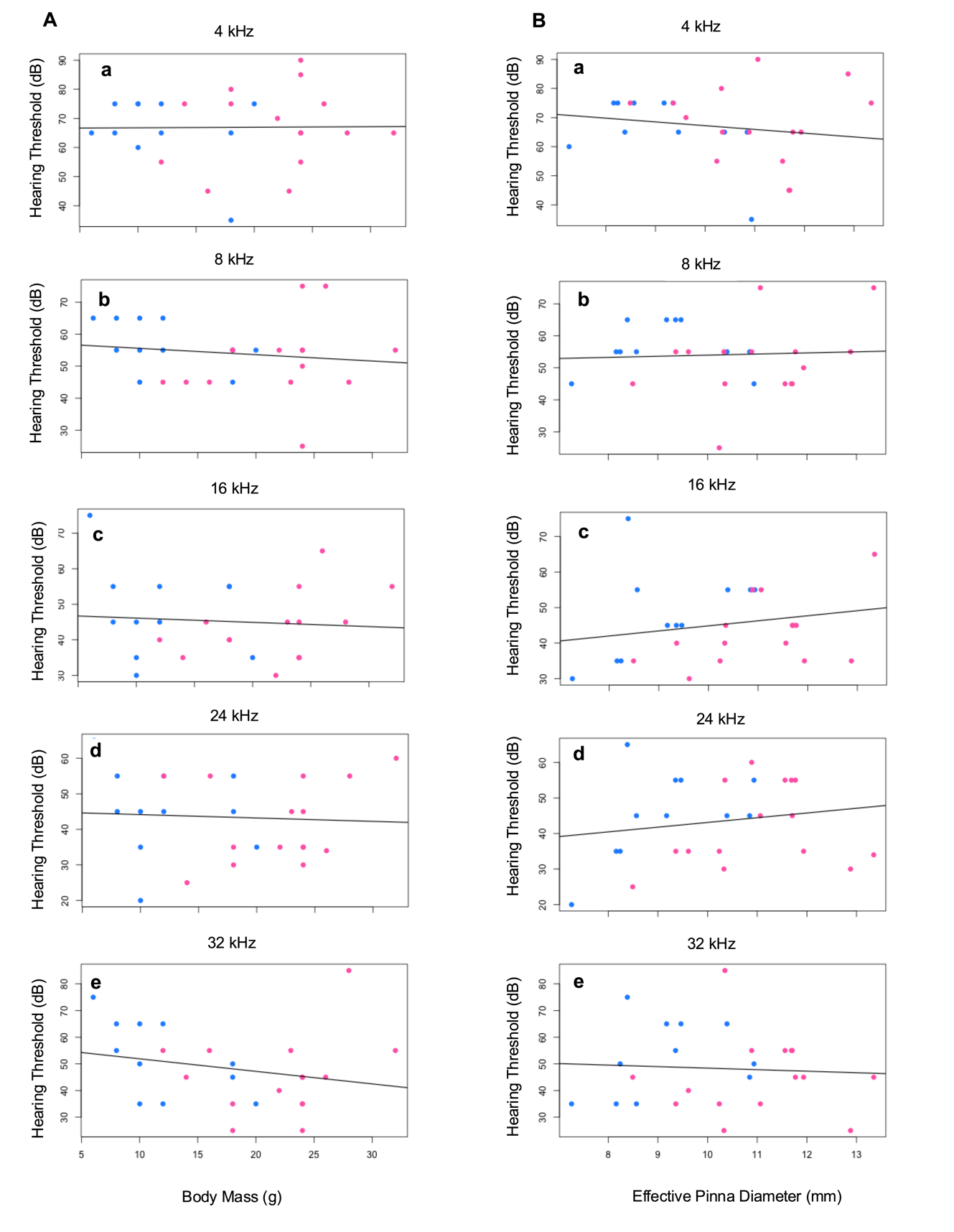

**Figure 5:** Scatter plots of frequency thresholds and morphological features (A = Body Mass, B = Effective Pinna Diameter) between species. Blue dots represent *P.* *maniculatus* and pink dots represent *P. leucopus*. No significant effects of body mass and effective pinna diameter were detected on best frequency hearing thresholds between species at 4, 8, 16, 24, and 32 kHz (All Pvalue < 0.05)

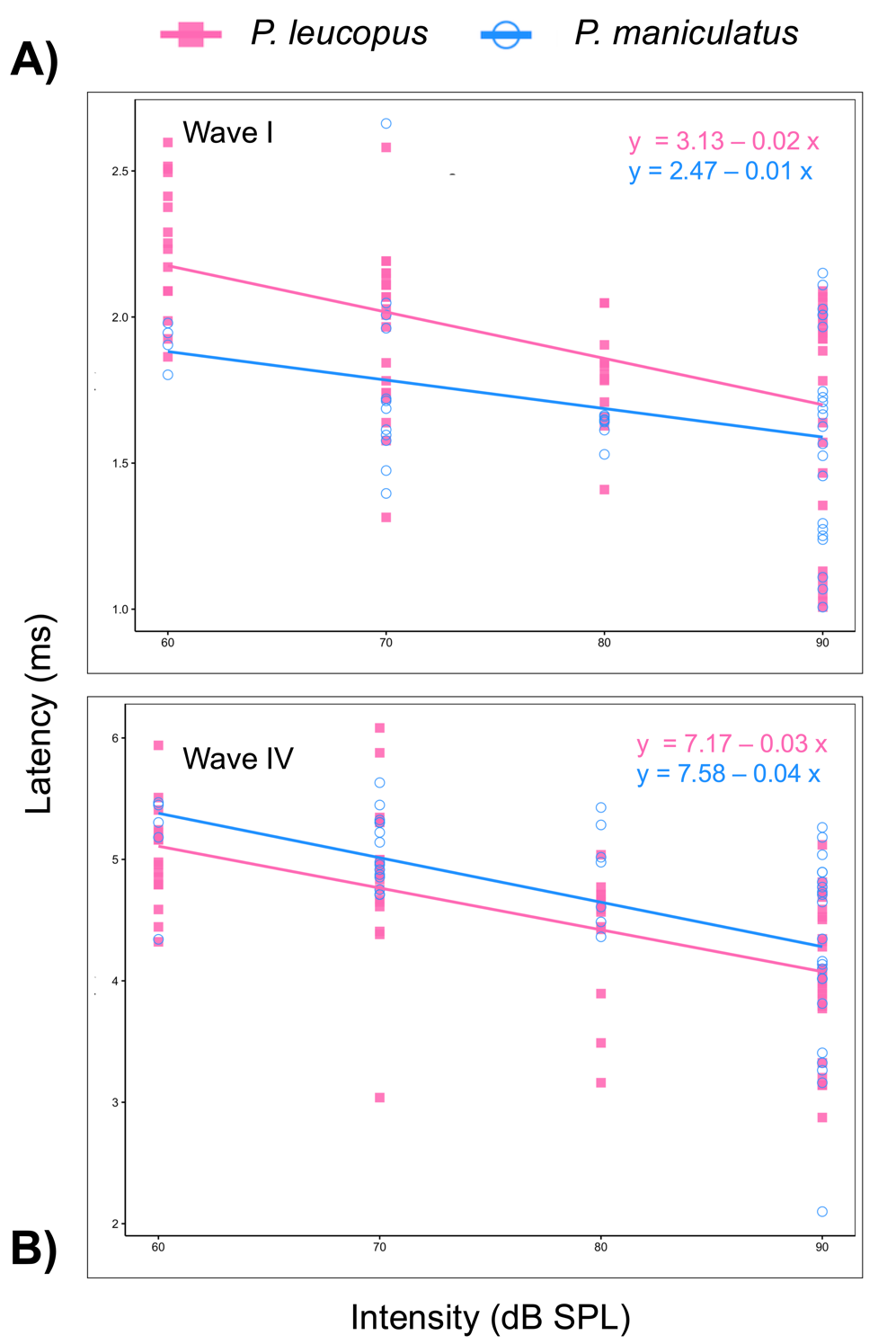

**Figure 6:** 6A shows latency data of wave I for both studied species with linear equation (slope). 6B shows latency data of wave IV for both studied species with linear equation (slope)

| Individuals | Species ID | Query Cover | Similarity | GenBank Accession number | Sex | Coordinates | Sequences |
| --- | --- | --- | --- | --- | --- | --- | --- |
| 1 | *P. maniculatus* | 98% | 98.32% | NC 039921.1 | Male | (36.19139; -96.9496) | (Barbour et al., 2019a) |
| 2 | *P. maniculatus* | 98% | 98.87% | NC 039921.1 | Female | (36.19139; -96.9496) | (Barbour et al., 2019a) |
| 3 | *P. maniculatus* | 99% | 98.75% | NC 039921.1 | Male | (36.19139; -96.9496) | (Barbour et al., 2019a) |
| 4 | *P. maniculatus* | 99% | 98.61% | NC 039921.1 | Male | (36.19139; -96.9496) | (Barbour et al., 2019a) |
| 5 | *P. maniculatus* | 96% | 99.14% | NC 039921.1 | Male | (36.19139; -96.9496) | (Barbour et al., 2019a) |
| 6 | *P. maniculatus* | 99% | 98.47% | NC 039921.1 | Male | (36.19139; -96.9496) | (Barbour et al., 2019a) |
| 7 | *P. maniculatus* | 97% | 99.15% | NC 039921.1 | Male | (36.19139; -96.9496) | (Barbour et al., 2019a) |
| 8 | *P. maniculatus* | 96% | 98.87% | NC 039921.1 | Male | (36.19139; -96.9496) | (Barbour et al., 2019a) |
| 9 | *P. maniculatus* | 98% | 98.52% | NC 039921.1 | Male | (36.19139; -96.9496) | (Barbour et al., 2019a) |
| 10 | *P. maniculatus* | 98% | 96.08% | NC 039921.1 | Female | (35.04399; -95.4719) | (Barbour et al., 2019a) |
| 11 | *P. maniculatus* | 98% | 94.02% | NC 039921.1 | Male | (35.04366; -95.4881) | (Barbour et al., 2019a) |
| 12 | *P. leucopus* | 98% | 98.32% | MH256659.1 | Male | (36.10736; -97.2285) | (Barbour et al., 2019b) |
| 13 | *P. leucopus* | 96% | 98.72% | MH256659.1 | Female | (36.10736; -97.2285) | (Barbour et al., 2019b) |
| 14 | *P. leucopus* | 99% | 98.06% | MH256659.1 | Female | (36.10736; -97.2285) | (Barbour et al., 2019b) |
| 15 | *P. leucopus* | 96% | 98.30% | MH256659.1 | Male | (36.10736; -97.2285) | (Barbour et al., 2019b) |
| 16 | *P. leucopus* | 98% | 98.32% | MH256659.1 | Female | (36.11208; -97.2285) | (Barbour et al., 2019b) |
| 17 | *P. leucopus* | 97% | 98.18% | MH256659.1 | Male | (36.11208; -97.2285) | (Barbour et al., 2019b) |
| 18 | *P. leucopus* | 96% | 98.57% | MH256659.1 | Male | (36.10736; -97.2285) | (Barbour et al., 2019b) |
| 19 | *P. leucopus* | 97 % | 98.32% | MH256659.1 | Female | (36.11208; -97.2285) | (Barbour et al., 2019b) |
| 20 | *P. leucopus* | 97% | 98.18% | MH256659.1 | Male | (36.10736; -97.2285) | (Barbour et al., 2019b) |
| 21 | *P. leucopus* | 97% | 98.18% | MH256659.1 | Male | (36.10736; -97.2285) | (Barbour et al., 2019b) |
| 22 | *P. leucopus* | 96% | 98.72% | MH256659.1 | Male | (35.89808; -99.7401) | (Barbour et al., 2019b) |
| 23 | *P. leucopus* | 97% | 98.04% | MH256659.1 | Female | (36.10736; -97.2285) | (Barbour et al., 2019b) |
| 24 | *P. leucopus* | 97% | 98.45% | MH256659.1 | Male | (35.87615; - 99.649) | (Barbour et al., 2019b) |
| 25 | *P. leucopus* | 96% | 98.71% | MH256659.1 | Female | (35.89383; -99.6614) | (Barbour et al., 2019b) |
| 26 | *P. leucopus* | 97% | 97.63% | MH256659.1 | Male | (35.89383; -99.6614) | (Barbour et al., 2019b) |

Table 1: Provenance of *P. leucopus* and *P. maniculatus* samples. Query cover represent percent overlap and similarity represent percent similarity when compared to other sequencing on the GenBank data
